# Intracellular Lactate Dynamics in *Drosophila* Glutamatergic Neurons

**DOI:** 10.1101/2024.02.26.582095

**Authors:** Matthew S. Price, Elham Rastegari, Richa Gupta, Katie Vo, Travis I. Moore, Kartik Venkatachalam

## Abstract

Rates of lactate production and consumption reflect the metabolic state of many cell types, including neurons. Here, we investigate the effects of nutrient deprivation on lactate dynamics in *Drosophila* glutamatergic neurons by leveraging the limiting effects of the diffusion barrier surrounding cells in culture. We found that neurons constitutively consume lactate when availability of trehalose, the glucose disaccharide preferred by insects, is limited by the diffusion barrier. Acute mechanical disruption of the barrier reduced this reliance on lactate. Through kinetic modeling and experimental validation, we demonstrate that neuronal lactate consumption rates correlate inversely with their mitochondrial density. Further, we found that lactate levels in neurons exhibited temporal correlations that allowed prediction of cytosolic lactate dynamics after the disruption of the diffusion barrier from pre-perturbation lactate fluctuations. Collectively, our findings reveal the influence of diffusion barriers on neuronal metabolic preferences, and demonstrate the existence of temporal correlations between lactate dynamics under conditions of nutrient deprivation and those evoked by the subsequent restoration of nutrient availability.

## INTRODUCTION

The advent of genetically-encoded sensors of energy-related metabolites, such as ATP, glucose, pyruvate, lactate, and the NAD^+^/NADH ratio, has revolutionized our understanding of the relationship between neuronal activity and bioenergetics^1^. By enabling real-time assessment of metabolic dynamics at cellular resolution, these sensors have elucidated how neuronal activity affects ATP production and revealed mechanisms underlying the metabolic burden of neuronal excitability, Ca^2+^ extrusion, and synaptic transmission^2–7^. Using these tools, pioneering work in *Drosophila* has revealed strong correlations between activity and metabolism in neuronal circuits of freely behaving animals^1,8^. These studies demonstrate that neurons allocate metabolic resources based on historical activity patterns, and that the metabolic states of interconnected neurons are correlated^8–11^.

Lactate is a metabolite that stands at the crossroads of glycolysis, mitochondrial ATP production, and cellular redox state. Its production depends on the reduction of glycolysis-derived pyruvate by lactate dehydrogenase (LDH)^12,13^. In energetically demanding cells such as neurons, lactate oxidation is a glucose-sparing source of pyruvate, which is taken up into the mitochondria to power the tricarboxylic acid (TCA) cycle and oxidative phosphorylation (OXPHOS)^14,15^. In response to stimulation, mouse hippocampal neurons augment glycolysis, which temporarily exceeds OXPHOS, leading to the accumulation of lactate^3^. Neuronally-generated lactate can either be extruded or subsequently oxidized to pyruvate for the sustenance of mitochondrial ATP synthesis. In addition to this cell-autonomous axis of lactate production, glia can also supply neurons with lactate as a substrate for energy production, especially during periods of heightened neuronal activity and computational load^15–19^. Oxidation of glia-derived lactate within neurons is necessary for the regulation of neuronal excitability, plasticity, and viability^15,17,19,20^. Notably, direct application of lactate, rather than glucose, increases neuronal spiking activity in rodent cortical neurons^21^. Examinations of lactate flux (i.e., changes in lactate levels over time) can also provide insights into the metabolic state of neurons. For instance, preferential use of lactate as a metabolic substrate reflects dependance on OXPHOS for energy production^21^. Since the cytosolic NAD^+^/NADH ratio is a readout of the directionality of LDH activity (pyruvate-to-lactate versus lactate-to-pyruvate)^14,17^, changes in lactate concentration also reflect the cellular redox balance^12^.

In this study, our objective was to examine lactate dynamics at cellular resolution in *Drosophila* neurons. Using the FRET-based lactate biosensor, Laconic^22^, we found that dissociated *Drosophila* glutamatergic neurons are constitutive lactate consumers, albeit with notable cell-to-cell variability. Interestingly, the imaged neurons consumed lactate even in the presence of adequate levels of trehalose — the preferred sugar substrate in *Drosophila* and other insects^23–25^. Further analysis revealed that this preference for lactate consumption in the presence of trehalose reflected the formation of a diffusion barrier around the neurons referred to as the “unstirred layer”^26–29^. We found that mechanical disruption of the unstirred layer prevented the limiting effects, and thus, reduced the rates of lactate consumption. The effects of buffer mixing on the unstirred layer and the rates of lactate consumption were augmented by the inclusion of lactate and trehalose in the mixing buffer.

While the limiting effects of the unstirred layer on metabolite and oxygen availability presents a technical challenge in some contexts, we have leveraged this phenomenon as a biologically-meaningful opportunity for studying neuronal responses to nutrient deprivation. For instance, the limiting effects of the unstirred layer on trehalose availability indicates that *Drosophila* glutamatergic neurons can simultaneously oxidize both lactate and glucose (derived from trehalose) to generate ATP. Kinetic modeling and experimental validation suggested that variations in cellular oxygen consumption, reflected in mitochondrial density, underlie the observed heterogeneity in lactate consumption rates under unstirred layer conditions. These findings provide insights into the general determinants of glucose versus lactate oxidation in fly neurons. Further, mechanical disruption of the unstirred layer revealed that fluctuations in lactate levels showed strong temporal correlations that enabled accurate prediction of postmixing responses from premixing lactate dynamics. The addition of ectopic lactate and metabolizable sugars, disrupted these temporal patterns without affecting steady-state lactate levels. Therefore, we harnessed the nutrient-limiting effects of the unstirred layer to demonstrate that neuronal metabolic responses appear to exist along a spectrum, from highly predictable responses determined by preexisting states to more complex responses where external metabolites can reorganize intrinsic patterns.

## RESULTS

### Dissociated *Drosophila* glutamatergic neurons exhibit dynamic changes in cytosolic lactate levels

We investigated cytosolic lactate dynamics using Laconic — a FRET-based lactate biosensor comprised of the lactate binding domain of the bacterial transcription factor, LldR, flanked by mTFP and Venus^22^. In this system, proximity of the fluorophores in low cytosolic [lactate] results in constitutive mTFP–Venus FRET, which is attenuated by an increase in cytosolic [lactate] ^22,30^. Therefore, changes in cytosolic [lactate] are represented by the inverse of mTFP–Venus FRET efficiency, which we refer to as the Laconic ratio^22,30^. We expressed *UAS-Laconic*^31^ under the control of the *d42-GAL4* glutamatergic/motor neuron driver^32,33^ (*d42>Laconic*) and performed live imaging on neurons dissociated from 3^rd^ instar larval brains (Figure 1A) in hemolymph-like buffer (HL3, pH 7.2) containing trehalose and sucrose, but without glucose or lactate^34^. Normalization of the Laconic ratios to starting values revealed that the ratios dropped steadily in ∼75% of dissociated glutamatergic neurons (Figure 1B). Over six minutes, average Laconic ratios in the neuronal population dropped ∼10% below baseline (Figures 1B and 1D). Analysis of individual neurons revealed exponential decay kinetics with rate constants (k) significantly greater than zero (Figure 1E), demonstrating reductions in cytosolic [lactate] at rest.

**Figure 1.**
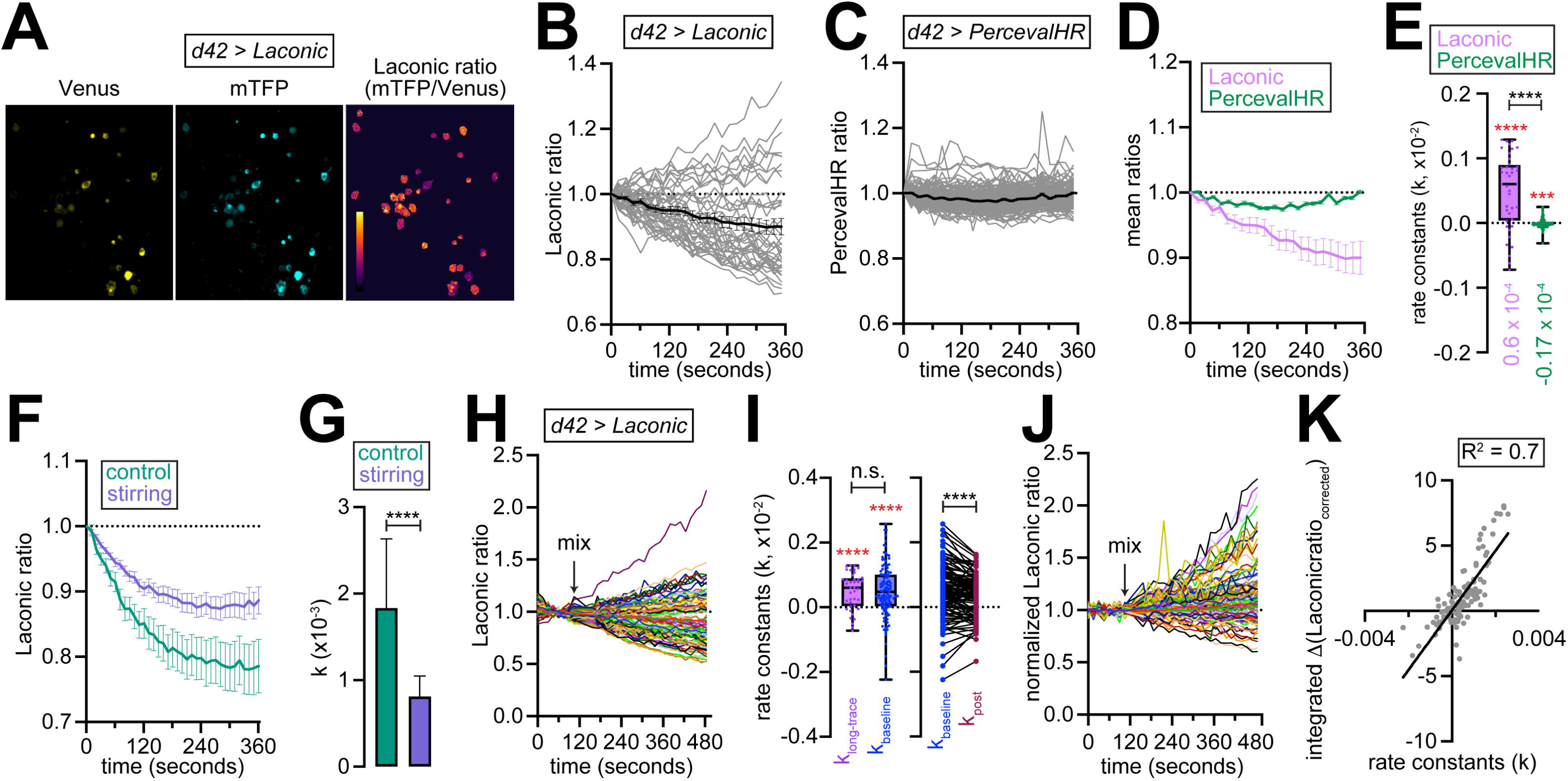
Impact of the unstirred layer on neuronal lactate metabolism. **(A)** Representative images of neurons dissociated from brains of *Drosophila* larvae expressing *UAS-Laconic* under the control of *d42-GAL4* glutamatergic/motor neuron driver (*d42>Laconic* neurons). Venus and mTFP intensities and the ratio of the intensities (Laconic ratio) are shown. **(B)** Traces showing Laconic ratios in *d42>Laconic* neurons. Each trace is normalized to the starting value of that trace. Responses of individual neurons are shown in grey. Black line with error bars shows mean ± SEM of the population of neurons. **(C)** Traces showing PercevalHR ratios in *d42>PercevalHR* neurons. Each trace is normalized to the starting value of that trace. Responses of individual neurons are shown in grey. Black line shows mean ± SEM of the population. **(D)** Traces showing mean ± SEM normalized Laconic and PercevalHR ratios from panels **(B)** and **(C)**, respectively. **(E)** Boxplots showing rate constants of exponential fits applied to Laconic and PercevalHR traces from panels **(B)** and **(C)**, respectively. Dots represent individual values. ****, *P* < 0.0001, ***, *P* < 0.001. Red asterisks represent results of one-sample Wilcoxon test to examine difference from a median value of 0. Black asterisks are from Mann Whitney tests. **(F)** Traces showing mean ± SEM Laconic ratios normalized to the starting values in control neurons or in neurons subjected to stirring. **(G)** Bar graph showing the rate constants of exponential fits applied to Laconic traces shown in panel **(F)**. Values represent median ± 95% CI. ****, *P* < 0.0001, Mann Whitney tests. **(H)** Traces showing Laconic ratios in *d42>Laconic* neurons. Arrow represents time of buffer mixing. **(I)** *Left*, boxplot showing rate constants of exponential fits applied to the indicated traces. Dots represent individual values. *Right*, rate constants before and after buffer mixing. ****, *P* < 0.0001, n.s., not significant. Red asterisks represent results of one-sample Wilcoxon test to examine difference from a median value of 0. Black asterisks are from Mann Whitney tests. **(J)** Traces showing normalized Laconic ratios after pH correction. Arrow represents time of buffer mixing. **(K)** Scatterplot of the relationship between integrated changes in Laconic ratios and premixing rate constants. Regression line is shown.

### Relationship between lactate levels and energy status in dissociated Drosophila glutamatergic neurons

Lactate is a glucose-sparing source of pyruvate. Via the combined activities of the TCA and OXPHOS, lactate-derived pyruvate is used for the generation of ATP^12–15^. To determine whether the drop in cytosolic [lactate] reflected its utilization for supporting cellular bioenergetics, we monitored ATP/ADP ratios using the fluorescent reporter, PercevalHR^35–37^, in glutamatergic/motor neurons (*d42>PercevalHR*). Despite the continuous decrease in cytosolic [lactate], ATP/ADP ratios remained relatively stable, showing only a transient initial decline before returning to baseline by six minutes (Figures 1C-1D). When fitted to the exponential decay function, the rate constants of the changes in PercevalHR ratios were slightly negative and significantly lower than those for Laconic decay (Figure 1E). These patterns of lactate and ATP/ADP flux were consistent with dissociated neurons consuming lactate to maintain ATP levels.

### The “unstirred layer” and its impact neuronal lactate metabolism

Trehalose is the preferred sugar source in *Drosophila*^23–25^. Trehalose utilization for energy production in fly cells is initiated by trehalase-mediated breakdown of the disaccharide into glucose monomers, which are then oxidized during glycolysis^23^. The reaction catalyzed by the glycolytic enzyme, glyceraldehyde-3-phosphate dehydrogenase (GAPDH), decreases the cytosolic NAD^+^/NADH ratio and prompts LDH to convert pyruvate to lactate in order to restore NAD^+^ levels for the continuation of glycolysis^38^. Conversely, when glucose availability is limited and NAD^+^/NADH ratios rise, LDH operates in reverse, thereby oxidizing lactate to pyruvate while generating NADH. This metabolic toggle between glycolysis and lactate oxidation is largely governed by glucose availability.

Given this well-established metabolic framework, we asked why dissociated neurons would be consuming lactate in the presence of abundant trehalose (5 mM). One possible explanation for this observation was a diffusion barrier introduced by the formation of a stagnant, unstirred layer around the neurons. A phenomenon first described in the 1960s, the unstirred layer is formed as a result of solute and osmotic flow across cell membranes resulting in localized concentration changes adjacent to the membrane^26–29^. Discontinuity in diffusion rates introduced by the unstirred layer can reduce solute concentration within the layer. Of pertinence to cellular metabolism, the unstirred layer has been shown to limit the uptake of nutrients as well as oxygen into different cell types, including neurons^28,39^. Although an unstirred layer with no internal convective mixing will form adjacent to cell membrane even with perfect stirring in the bulk solution^27^, experimental evidence has shown that stirring thins the unstirred layer^29^.

We reasoned that that if an unstirred layer formed around the imaged neurons were limiting the continued uptake of nutrients (e.g., trehalose), those neurons would be forced to rely on lactate oxidation for energy production. If so, stirring the bulk solution might counteract these effects, as indicated previously^29^. To test this idea, we introduced gentle stirring into the bulk solution using a magnetic stir-bar. As would be expected from thinning of the unstirred layer^29^, stirring significantly lowered the rates of cytosolic lactate consumption in neurons (Figures S1A and 1F-1G). To examine further the effects of buffer mixing on the kinetics of lactate consumption, we compared the kinetics of lactate consumption before and after the rapid application of 20 μL HL3 to the existing 100 μL solution (Figure 1H). While premixing rate constants matched those observed in 6-minute recordings, mixing led to significant reduction in the rates of lactate consumption in those neurons (Figures 1H-1I). Since the composition of the newly added buffer was the same as that to which the cells were already exposed, these outcomes are likely the result of the acute mechanical disruption of the unstirred layer.

To account for the potential effects of cytosolic pH on Laconic measurements, we performed simultaneous pH measurements using pHrodo-AM. We utilized an empirical pH correction protocol that first normalized the Laconic ratios to the exponential fits of the premixing baselines (Figure S1B). By doing so, average pre-mixing baselines were set to 1, and reductions in the rates of lactate consumption after mixing appeared as upward deflections of the ratios (Figure S1B). Cytosolic alkalinization induced by application of 10 mM NH_4_Cl led to acute decreases in both the normalized Laconic ratios and pHrodo fluorescence (Figure S1C). Upon removal of NH_4_Cl, both pHrodo fluorescence and normalized Laconic ratios exhibited transient overshoots reflecting the compensatory response of neurons to cytosolic alkalinization (Figure S1C). Deviations of the pHrodo and Laconic signals from baseline (ΔpHrodo and ΔLaconic ratio, respectively) exhibited linear relationships (Figure S1D), which were used to empirically correct for the changes in Laconic ratios that stem solely from changes in cytosolic pH. This approach corrected the effects of cytosolic pH changes on the normalized Laconic ratios (Figure S1E). By applying NH_4_Cl pulses at the end of each recording, we obtained pH-corrected normalized Laconic responses (Figure 1J). These traces revealed that the integrated changes in normalized Laconic ratios exhibited linear relationship with premixing rate constants (Figure 1K), strengthening our conclusion that mixing affected neuronal lactate metabolism by disrupting the unstirred layer.

### Simple kinetic models capture the effects of buffer mixing on neuronal lactate consumption

To understand the mechanistic basis of the effects of buffer mixing on neuronal lactate consumption, we developed a simple kinetic model tracking the dynamics of [trehalose], [lactate], [ATP], and [O_2_]. This “toy model” focused on assessing qualitative behavior of the system, and incorporated variable ATP demand and O_2_ consumption rates (k_oxygen_) (Figure 2A). Our model had the following features: ***(1)*** ATP synthesis was calibrated to match demand, as has been shown experimentally^6^; ***(2)*** lactate production was coupled to glycolysis (simulating LDH-mediated NAD^+^ replenishment); and ***(3)*** it switched to lactate oxidation for ATP production as [trehalose] decreased. Under conditions mimicking high ATP demand and O_2_ consumption rates, the model predicted [lactate] accumulation before mixing followed by further increases after mixing (Figure 2B, left — grey box represents the imaged period). This pattern reflected preferential trehalose consumption and lactate production under low [O_2_] conditions. With lower ATP demand and O_2_ consumption rates, adequate oxygen availability permitted lactate consumption followed by rapid [lactate] increases after mixing (Figure 2B, right). Upon normalization to premixing baselines, we observed variable lactate levels in response to different ATP demands and O_2_ consumption rates (Figure 2C). However, these predicted variations in [lactate] showed two key discrepancies when compared to experimental data. First, mixing led to universal increases in normalized [lactate] (Figure 2C), which was contrary to the experimental observation of both increases and decreases in normalized Laconic ratios after mixing (Figure 1J). Second, integrated [lactate] changes exhibited a J-shaped relationship with premixing rate constants (Figure 2D), rather than the experimentally observed linear relationship (Figure 1K).

**Figure 2.**
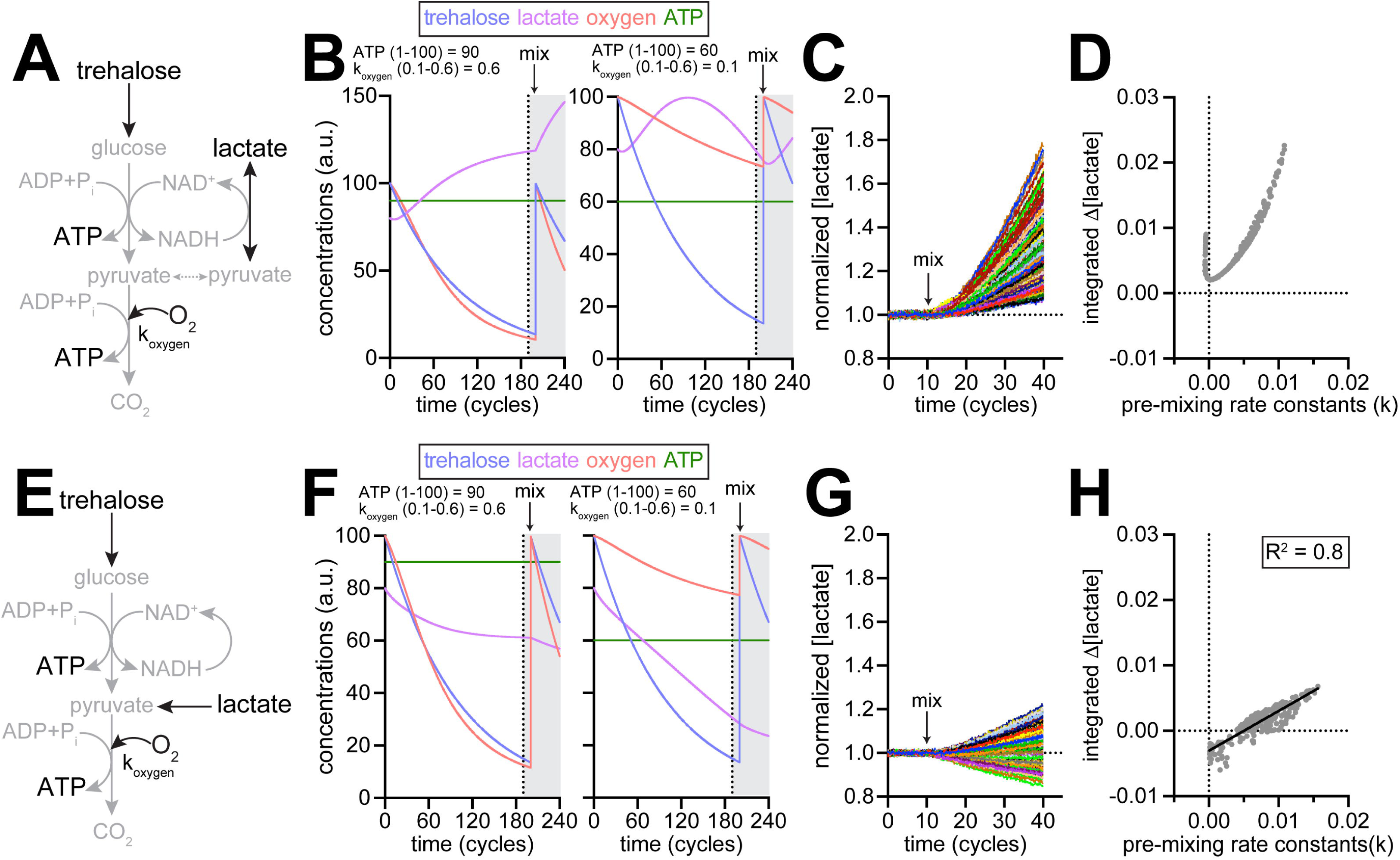
Simple kinetic models of the effects of buffer mixing on neuronal metabolite levels. (**A and E**) Models of metabolite fluxes. Lactate production is coupled to glucose oxidation in **(A)** but not in **(E)**. Metabolites indicated in black are explicitly modeled whereas the metabolites in grey are implicit. **(B)** Representative predictions of metabolite concentration dynamics based on the model shown in **(A)**. Arbitrary values of ATP demand and rates of oxygen consumption (k_oxygen_) are indicated above the traces. Grey box represents the epoch equivalent to the imaging period. Arrow represents point in time when [trehalose] of [O_2_] are restored to starting values, as would occur in response to buffer mixing. **(C)** Normalized predicted lactate responses from 300 runs of the model shown in **(A)** with randomly selected values for ATP demand and k_oxygen_ from ranges of 1-100 and 0.1-0.6, respectively. **(D)** Scatterplot of the relationship between predicted integrated changes in lactate levels and predicted premixing rate constants. **(F)** Same as **(B)** but based on the model shown in **(E)**. **(G)** Same as **(C)** but based on the model shown in **(E)**. **(H)** Same as **(D)** but based on the model shown in **(E)**. Regression line is shown.

To resolve these discrepancies, we refined the model by uncoupling lactate production from glycolysis (Figure 2E). This modification reflected the possibility that oxidation of NADH to NAD^+^ to facilitate the continued activity of GAPDH could involve a dehydrogenase other than LDH (e.g., glycerol-3-phosphate dehydrogenase, as suggested in flies and other organisms^12,38,40^). The refined model transformed cells into pure lactate consumers from being both producers and consumers of lactate, while retaining the repressive effect of [trehalose] on lactate consumption rates. In this new model, high ATP demand and O_2_ consumption were associated with minimal premixing [lactate] changes, followed by increased lactate consumption after mixing (Figure 2F, left). Lower ATP demand and O_2_ consumption rates predicted rates of decline in premixing [lactate], followed by reduced consumption rates after mixing (Figure 2F, right). These refined predictions better matched experimental observations. Normalization to premixing baselines revealed bidirectional changes in [lactate] after mixing (Figure 2G), which was consistent with experimental data (Figure 1J). Furthermore, integrated [lactate] changes exhibited a linear relationship with premixing rate constants (Figure 2H), which also matched the experimental observations (Figure 1K). These results demonstrate that decoupling lactate production from glycolysis, while maintaining trehalose-dependent regulation of lactate consumption, successfully reproduces the experimentally observed effects of buffer mixing on neuronal lactate consumption.

### Direct measurements of cytosolic glucose validate the effects of the unstirred layer

Next, we asked whether the limiting effects of the unstirred layer manifested as decreasing cytosolic glucose levels. Using a FRET-based glucose sensor, FLII12Pglu-700μδ6^41,42^, we monitored cytosolic [glucose] in dissociated glutamatergic neurons. Glucose measurements revealed patterns similar to those observed with lactate. FLII12Pglu-700μδ6 ratios exhibited constitutive exponential decreases with time, and buffer mixing significantly reduced these decay rates (Figures S2A-S2B). This observation was noteworthy given that the HL3 buffer contained no glucose. Therefore, the slower decline in [glucose] after mixing likely reflects enhanced rates of glucose production stemming from increased trehalose availability, consistent with mechanical disruption of the unstirred layer increasing trehalose accessibility, and therefore, glucose levels within the neurons.

### Determinants of the rates of neuronal lactate consumption

Our experimental observations revealed heterogenous lactate consumption rates in neurons owing to the limiting effects of unstirred layers. A kinetic model successfully reproduced this variability by incorporating different ATP demands and oxygen consumption rates. Direct comparisons between ATP demand, rates of oxygen consumption and premixing rates of lactate consumption, as predicted by the kinetic model, revealed that the highest rates lactate consumption corresponded to the lowest rates of oxygen consumption and ATP demand (Figure 3A). Rates of lactate consumption exhibited significant negative correlation with both the rates of oxygen consumption and ATP demand, with the negative correlation being particularly strong with the rates of oxygen consumption (Figure 3B). This stronger inverse correlation with oxygen consumption rates suggested that the cellular capacity for oxidative metabolism might be a determinant of lactate utilization kinetics under conditions of insufficient trehalose availability.

**Figure 3.**
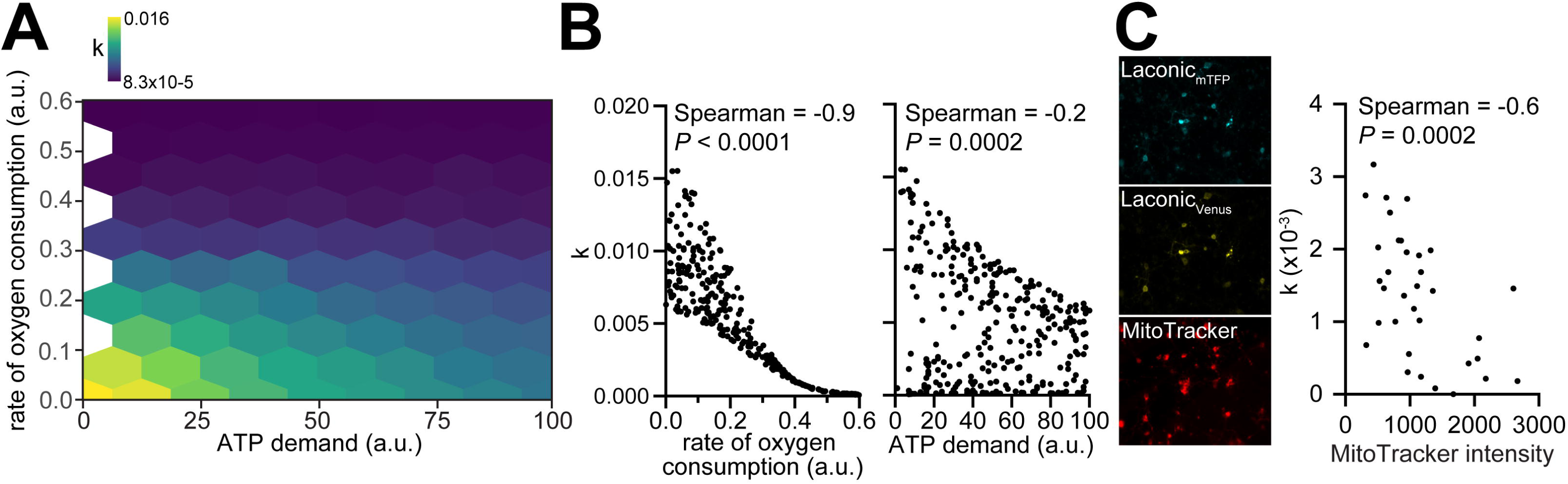
Determinants of the rates of neuronal lactate consumption. **(A)** Heatmap showing the relationships of O_2_ consumption rates and ATP demand with predicted rate constants of lactate decline derived from model shown in Figure 2E. **(B)** Scatterplots showing the correlation of predicted lactate consumption rates with the indicated parameters. **(C)** *Left,* representative confocal images of Laconic and MitoTracker in *d42>Laconic* neurons. *Right*, scatterplot showing the correlation between experimentally-determined lactate consumption rates and MitoTracker intensities in individual neurons.

To experimentally investigate these predictions, we probed the relationship between the rates of lactate consumption and mitochondrial density — a proxy for oxygen consumption capacity. By loading *d42>Laconic* neurons with MitoTracker, which accumulates in active mitochondria to an extent proportional to their membrane potential, we determined both the rates of lactate consumption and the content of functional mitochondria in the same neurons (Figure 3C). This analysis revealed a significant negative correlation between the rates of lactate consumption and MitoTracker intensity (Figure 3C), which matched our predictions. Thus, neuronal mitochondrial density, which directly influence oxygen consumption rates, correlates inversely with the rates of lactate consumption when trehalose becomes limiting.

Next, we assessed the potential involvement of other factors that could influence the rates of lactate consumption in the imaged neurons. Since only live neurons would consume lactate for ATP production, low rates of constitutive lactate consumption in some ROIs could indicate cell death. We assessed the viability of neurons on the dishes using the rhodamine-conjugated, membrane-impermeable LIVE/DEAD dye that enters cells only if the plasma membrane is compromised as a result of cell death. This analysis revealed that <1% (median value, ∼0.1%) of all the cells on a dish were stained by the LIVE/DEAD dye (Figure S3A). Permeabilization of the plasma membrane by application of 0.1% saponin led to the ∼10-fold increase in dye uptake (Figure S3A), indicating that the plasma membranes of live *Drosophila* brain cells on the dish are appropriate barriers to dye uptake. Therefore, cell death is a negligible factor in the variability of lactate consumption in the imaged neurons.

Since glia supply neurons with lactate as a substrate for energy production in the intact brain^15–19^, we also probed for the presence of glial cells in the vicinity of *d42>Laconic* neurons. After live imaging of Laconic ratios, we fixed several dishes containing *d42>Laconic* neurons and stained for glial nuclei using antibodies against the glial transcription factor, Repo^43^. None of the cells expressing *Laconic* showed immunoreactivity towards anti-Repo antibodies, indicating that *d42-GAL4* did not drive expression in glial cells, at least the ones plated on dishes. Further, each field of view was comprised of a significantly smaller fraction of glial cells relative to the fraction of glutamatergic neurons expressing *Laconic* (median fractions, ∼1% and ∼7%, respectively) (Figure S3B). This imbalance argued against glial cells on the dishes being major contributors of lactate for neuronal consumption. In addition, median distance between the centers of the anti-Repo-positive glial cells and *Laconic*-expressing neurons was ∼26 μm, which significantly exceeded the glial and neuronal radii (median radii, 3.3 and 2.0 μm, respectively) (Figure S3C). These distances between glia and neurons plated on the dishes argues further against the likelihood of glia contributing significantly to the rates of lactate consumption in the imaged neurons. Thus, rates of lactate consumption in neurons under our experimental conditions are driven cell autonomously with little or no contribution from glial cells.

### Impact of metabolite and drug inclusion in the mixing buffer

Having established that mechanical disruption of the unstirred layer reduces lactate consumption, we asked how the inclusion of various metabolites and drugs in the mixing buffer might affect this response. We found that inclusion of 10 mM lactate in the mixing buffer significantly enhanced the integrated change in Laconic ratio (Figure 4A, compare black and light blue bars), indicating that direct lactate supplementation synergized with mixing-induced decrease in lactate consumption. Both trehalose and glucose (5 mM each) also augmented the lactate responses relative to controls (Figure 4A, compare black bar with navy and green bars, respectively), which suggests that inclusion of metabolizable sugars in the mixing buffer resulted in stronger repression of lactate consumption. Arguing against effects of osmolarity changes upon inclusion of trehalose or glucose, inclusion of iso-osmolar levels of the metabolically-inert sugar, mannitol (5 mM), did not alter the Laconic responses (Figure 4A, compare black and 1^st^ brown bars). Neither the OXPHOS inhibitor, oligomycin A (oligo A, 10 μM), nor the inhibitor of pyruvate–lactate conversion by LDH, oxamate (10 mM), had any additional effects on the mixing response (Figure 4A, compare black bar with ruby and orange bars, respectively).

**Figure 4.**
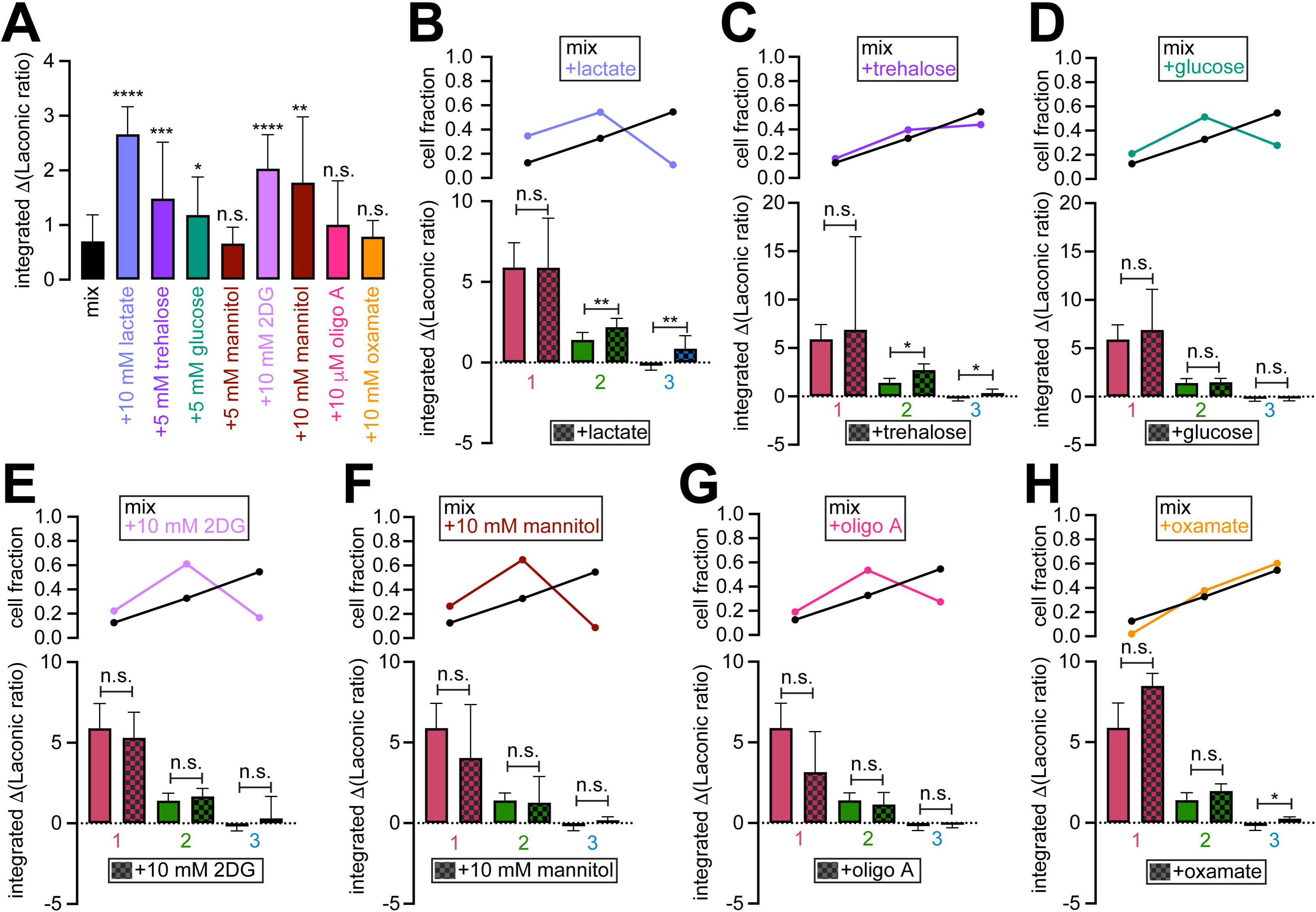
Impact of metabolite and drug inclusion in the mixing buffer on cytosolic Laconic ratios. **(A)** Bar graph showing integrated changes in the Laconic ratios in response to inclusion of the indicated concentrations of metabolites or drugs in the mixing buffer. Values represent median ± 95% CI. ****, *P* < 0.0001, ***, *P* < 0.001, **, *P* < 0.01, *, *P* < 0.05, n.s., not significant; pairwise Mann Whitney tests for differences from values obtained with mixing buffer without any metabolites or drugs. **(B-H)** *Top*, fraction of neurons belonging to the indicated clusters in response to the indicated treatments. *Bottom*, bar graph showing integrated changes in the Laconic ratios in the indicated clusters in response to inclusion of the indicated metabolite or drugs in the mixing buffer. Values represent median ± 95% CI. **, *P* < 0.01, *, *P* < 0.05, n.s., not significant; pairwise Mann Whitney tests for differences from values obtained with mixing buffer without any metabolites or drugs. Number of cells in clusters 1-3 and number of independent experiments are — mix (C1: 45, C2: 55, C3: 19, Expts: 10); 10 mM lactate (C1: 74, C2: 3, C3: 15, Expts: 6); 5 mM trehalose (C1: 44, C2: 5, C3: 44, Expts: 7); 5 mM glucose (C1: 14, C2: 26, C3: 75, Expts: 8); 5 mM mannitol (C1: 49, C2: 15, C3: 35, Expts: 8); 10 mM 2DG (C1: 18, C2: 52, C3: 20, Expts: 7); 10 mM mannitol (C1: 9, C2: 22, C3: 3, Expts: 3); 10 μM oligo A (C1: 16, C2: 45, C3: 23, Expts: 7); 10 mM oxamate (C1: 3, C2: 37, C3: 59, Expts: 7).

While the aforementioned findings made sense in the context of lactate metabolism, we obtained unexpected results upon inclusion of the non-hydrolyzable glucose analog and glycolysis inhibitor 2-deoxyglucose (2DG, 10 mM). 2DG appeared to enhance the effects of buffer mixing (Figure 4A, compare black with purple bars). Similarly unexpected was the enhancement of the mixing effect by 10 mM mannitol (Figure 4A, compare black with 2^nd^ brown bars). We reasoned that these puzzling results might stem from inherent heterogeneity in neuronal responses to the unstirred layer, as evident from the wide range of rate constants of premixing lactate consumption (from - 2.2*10^-3^ to 2.6*10^-3^). To ensure that we are comparing neurons with similar kinetics of lactate consumption, we used the “Jenks natural breaks classification” method (defines cluster boundaries by minimizing intra-cluster variance and maximizing inter-cluster variance, see *Methods*) to sort neurons into one of 3 internally-homogenous clusters on the basis of their rate constants. As evident in the case of mixing with HL3, clusters showed progressive ranges of rate constants from high to low (Figures S4A-S4B). Cluster 1 (highest rate constants) showed the largest changes in Laconic ratio, whereas clusters 2 and 3 showed progressively smaller changes (Figures S4C-S4D) — a reflection of the linear relationship between integrated Laconic changes and pre-mixing rate constants. Notably, neurons that shared membership in a particular cluster were often plated on different dishes or imaged on different days. Many neurons in a cluster did not even originate from the same animal. Therefore, patterns of neuronal lactate consumption stemmed from inherent, cell autonomous properties of the imaged neurons.

This clustering strategy enabled more precise “apple-to-apple” comparisons of responses within matching clusters and obviated the confounding effects of heterogenous lactate consumption rates. It revealed that inclusion of either lactate or trehalose augmented the effects of buffer mixing, but only in neurons that belonged to clusters 2 or 3 (bar-graphs, Figures 4B and 4C, respectively). In contrast, neither glucose nor 5 mM mannitol exerted significant influences on the integrated changes in Laconic ratio in any of the clusters (bar-graphs, Figures 4D and S4E, respectively). Therefore, the apparent stimulatory effect of glucose reflected the outcome of sampling bias, i.e., a greater fraction of sampled neurons belonged to clusters 1 or 2 and a smaller fraction of sampled neurons belonged to cluster 3 (cell fraction, Figure 4D). The apparent stimulatory effects of 2DG and 10 mM mannitol were also due to similar sampling bias, as intra-cluster comparisons revealed that neither drug had any influence on the effects of buffer mixing (Figures 4E and 4F, respectively). To determine whether 10 mM 2DG was appropriately inhibiting glycolysis, we evaluated the effect of the drug on neuronal ATP/ADP ratio. As expected, 2DG application led to an acute decrease in cytosolic ATP/ADP ratio (Figures S4F-S4G), which pointed to its inhibitory effect on glycolysis. That the inhibition of glycolysis did not have much of an influence on neuronal [lactate] suggests that a relatively small fraction of neuronal lactate is derived from local glycolysis. Lastly, we found that oligo A had no significant influences on the integrated changes in Laconic ratio in any of the clusters (bar-graphs, Figures 4G), while oxamate induced a slight but significant in the Laconic ratio in neurons belonging to cluster 3 (bar-graphs, Figures 4H).

### Premixing fluctuations in cytosolic lactate are correlated with neuronal responses to mixing-induced disruption of the unstirred layer

The analyses described above revealed a quantitative relationship between the premixing lactate consumption rates and the magnitudes of [lactate] changes upon disruption of the unstirred layer. Even after normalization, Laconic ratios exhibited distinct patterns of fluctuations around the baseline (e.g., Figure 1J), likely reflecting temporal variations in cellular metabolic demand, transporter activity etc. Because fluctuations in functional parameters can encode meaningful information about cell state and response capacity, we hypothesized that premixing variability in normalized Laconic ratios (i.e., fluctuations in baseline [lactate]) might predict the neurons’ responses to the subsequent disruption of the unstirred layer. To test this hypothesis, we developed a machine learning approach using a neural network with a single hidden layer that incorporated both dropout and L1/L2 regularization to prevent overfitting^44,45^ (Figure 5A). We trained this network using premixing Laconic ratio time series as input features and postmixing ratios as training labels (Figure 5B). The model’s predictive capacity was then tested on unseen premixing Laconic ratios (Figure 5B). Using 5-fold cross-validation, we demonstrated robust prediction accuracy for both training and test datasets, with predicted Laconic ratios closely matching experimental observations (Figures 5C and 5D, respectively). To validate that the model captured genuine biological relationships rather than spurious correlations, we scrambled the temporal order of postmixing Laconic ratios (i.e., scrambled the temporal order of the labels). This manipulation greatly reduced prediction accuracy (Figure 5E), confirming that the model learned authentic temporal patterns in the data. Mean square error (MSE) values for buffer mixing predictions were statistically indistinguishable between the training and test datasets (Figure 5F), which indicates successful generalization without overfitting. Scrambled response labels resulted in ∼6-fold higher MSE values (Figure 5F), supporting the existence of temporal relationships between pre- and postmixing Laconic ratios.

**Figure 5.**
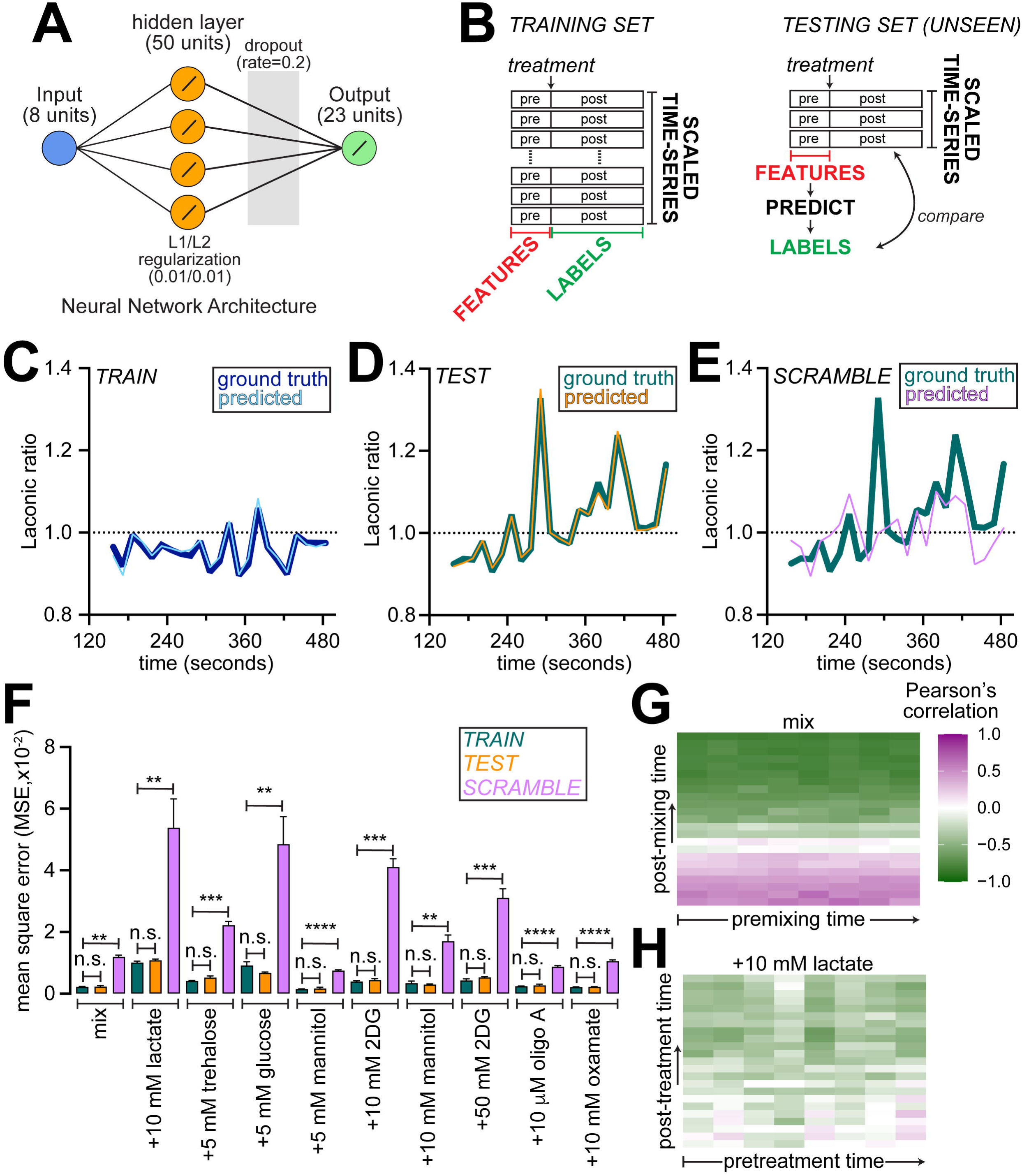
Correlations between premixing fluctuations in cytosolic lactate and the responses to disruption of the unstirred layer. **(A)** Schematic of the neural network architecture used for prediction of post-mixing Laconic ratios from premixing values. Linear activation functions used in each layer are indicated. **(B)** Schematic showing the training and testing protocols for the neural network designed to predict post-mixing Laconic ratios from premixing values. **(C)** Representative Laconic traces showing the ground truth and predicted post-mixing responses from traces used to train the model. **(D)** Representative Laconic traces showing the ground truth and predicted post-mixing responses from unseen (test) traces. **(E)** Representative Laconic traces showing the ground truth and predicted post-mixing responses from traces when the model was trained using labels with scrambled temporal order. **(F)** Bar graph showing the mean square error (MSE) values from comparison of the predicted values with the ground truth. Values represent mean ± SEM. ****, *P* < 0.0001, ***, *P* < 0.001, **, *P*<0.01%, n.s., not significant, t-tests or Mann Whitney tests depending on data normality. **(G)** Heatmaps showing Pearson’s correlation between the Laconic ratios of in all neurons at each premixing time (columns) and Laconic ratios in the same neurons at each post-mixing time (rows). **(H)** Same as **(G)** with inclusion of 10 mM lactate in the mixing buffer. Heatmap scale shown in panel **(G)** also applies to panel **(H)**.

This predictive capability of our model extended beyond buffer mixing. Comparable prediction accuracies were achieved for responses obtained by the inclusion of 5 mM mannitol, oligo A, and LDH inhibitor, oxamate, in the mixing buffer (Figure 5F). However, the inclusion of other metabolites and drugs in the mixing buffer partially disrupted the predictive relationship between the pre- and postmixing Laconic ratios. Higher MSE values were observed when the mixing buffer contained lactate, trehalose, glucose, 2DG, or 10 mM mannitol, although the model was still able to learn the relationship between premixing fluctuations and postmixing responses in all these cases (Figure 5F). Inclusion of lactate in the mixing buffer induced the greatest decline in model accuracy (Figure 5F), which suggested that uptake of lactate overrode the neurons’ initial response to the limiting effects of the unstirred layer.

Regardless of which premixing dataset was used for training, the model achieved similar prediction accuracies across all postmixing conditions (Figure S5A). This finding suggests the predictive capability of our model was robust, and pointed to the existence of fundamental temporal patterns in cellular lactate fluctuations that generalized across different conditions. In addition, temporal correlation analysis revealed that in response to buffer mixing or the inclusion of 5 mM mannitol in the mixing buffer, premixing Laconic ratios showed strong sequential correlations to postmixing ratios, transitioning from positive to negative correlation patterns over time (Figures 5G and S5B, respectively). This structured temporal relationship was greatly disrupted by the inclusion of lactate (Figure 5H), whereas inclusion of trehalose, glucose or 2DG in the mixing buffer led to partial disruption in the patterns of temporal correlations (Figures S5C-S5E). These data argue further that entry of some, though not all, ectopically added metabolites can either override or obscure the intrinsic rhythms of cytosolic lactate fluctuations.

## DISCUSSION

Our study reveals how the unstirred layer, a diffusion-limited boundary surrounding cells, can be leveraged to uncover properties of neuronal metabolism. By examining lactate dynamics in dissociated *Drosophila* neurons in unstirred layer conditions, we uncovered several insights into cellular metabolic regulation and heterogeneity. We demonstrated that *Drosophila* glutamatergic neurons are constitutive lactate consumers in unstirred layer conditions. Thus, local diffusion barriers can fundamentally alter cellular metabolic preferences. Since the effects of unstirred layer have also been observed *in vivo*, for instance in kidney tubules or during oxygenation of red blood cells^26,46,47^, it is possible that insect brains have evolved to constrain spatial distances between adjacent cell membranes in order to minimize the effects of such diffusion barriers.

Kinetic modeling revealed that by decoupling lactate production from glycolysis while maintaining trehalose-dependent regulation of lactate consumption, we successfully reproduced experimental observations. Thus, *Drosophila* glutamatergic neurons possess alternative (i.e., LDH independent) mechanisms for maintaining NAD^+^/NADH balance during glycolysis. Indeed, other dehydrogenases such as glycerol-3-phosphate dehydrogenase — part of the glycerol phosphate shuttle — have been shown to play roles in NAD^+^ replenishment during glycolysis^12,38,40^. This decoupling of lactate production from glycolysis also enables neurons to utilize glycolytically-generated pyruvate for TCA and OXPHOS-dependent mitochondrial ATP production. As such, these neurons simultaneously consume both trehalose and lactate, with the latter likely being supplied by local glial cells^15–19^. Such metabolic flexibility might be particularly important under conditions where trehalose and oxygen availability may be limited.

The inverse correlation between mitochondrial density and lactate consumption rates suggests that cells with lower oxidative capacity, and by extension, greater reliance on trehalose/glucose metabolism (i.e., neurons that are relatively more glycolytic), exhibit a greater propensity to switch to lactate metabolism in response to trehalose limitations. In other words, neurons that preferentially consume trehalose exhibit a more acute response to the limiting effects of that sugar. Neurons with higher oxidative capacity are likely better equipped to utilize metabolic substrates other than trehalose, thus making them less responsive to diffusion limitations. As demand-driven variations in neuronal metabolism correlate with cell-to-cell variations in firing properties^48^, it is also possible that the observed differences the extent of lactate consumption are related to the neurons’ firing possibilities. For instance, in both *Drosophila* and crayfish, motor neurons that exhibit tonic firing have greater mitochondrial density and oxidative activity than do the neurons with phasic (burst-like) firing patterns^31,49–51^. Future studies could be designed to ask phasic motor neurons, which have lower mitochondrial density that the tonic motor neurons, exhibit enhanced propensity for lactate consumption under conditions of trehalose limitation.

On the basis of premixing lactate consumption rates, we classified neurons into 3 subgroups of decreasing reliance on lactate. Classification into clusters ensured that all comparisons were made between groups of neurons with similar kinetics of lactate consumption. This analysis revealed that inclusion of 10 mM lactate in the mixing buffer led to increases in cytosolic lactate specifically in clusters with lower rates of lactate consumption (i.e., clusters 2 and 3 rather than cluster 1, Figure 4B). This observation stands to reason as neurons with inherently lower lactate consumption rates will be the ones that will naturally accumulate lactate. In neurons that are rapidly consuming lactate, ectopically added lactate will simply not have an opportunity to accumulate. Inclusion of trehalose in the mixing buffer also had similarly augmenting effects on cytosolic lactate in neurons belonging to clusters 2 and 3, indicating that trehalose supplementation resulted in significantly stronger effects on neurons with inherently slower lactate consumption rates. None of the other metabolites or drugs (including 2DG) had any appreciable influence on the effects of buffer mixing on the rates of lactate consumption. The observation that 2DG-mediated inhibition of glycolysis had no appreciable effect on neuronal lactate metabolism agrees further with the notion that neuronal lactate metabolism is decoupled from glycolysis.

Perhaps most intriguingly, we discovered that premixing fluctuations in cellular lactate levels contain predictive information about how neurons will respond to unstirred layer disruption. The ability of machine learning models to accurately predict postmixing responses from premixing dynamics suggested the existence of cell-intrinsic temporal correlations, as demonstrated by patterns of positive and negative correlations between pre- and postmixing Laconic ratios. Disruption of these temporal correlations by certain metabolites, particularly lactate, reveals how external factors can override or reorganize these intrinsic metabolic patterns. Notably, some compounds like 2DG disrupted temporal correlations without causing a net change in steady-state lactate levels. This outcome points to forms of metabolic reorganization that selectively affect the temporal dynamics of the fluctuations in metabolite concentrations. Thus, neuronal responses to the disruption of the unstirred layer exist along a spectrum — from highly predictable responses determined by pre-existing fluctuations to more complex responses where external metabolites can reorganize intrinsic patterns. Future studies could be directed towards understanding whether these differences in metabolic states align with established motor neuron subtypes^52–54^.

### Limitations of the study and future directions

Alterations in cytosolic [lactate] that we describe in this study occurred in neurons dissociated from the brains of 3^rd^ instar larvae. Dissociation of brains was needed to ensure that the applied metabolites or drugs had direct access to neurons without having to traverse the glial blood brain barrier. Changes in neuronal [lactate] would likely be different in the intact brain where neurons receive trophic and metabolic support from glia^16,17^. To overcome this limitation in future studies, we hope to examine neuronal [lactate] in intact brains, and the relationship between this metabolite and neuronal activity. These studies would nicely complement prior work focused on the relationships between activity and the levels of pyruvate and ATP in brains of live flies^8^.

While our studies have revealed that *d42-GAL4*-labeled neurons exhibit variable patterns of lactate flux stemming from differences in mitochondrial density, whether these neurons constitute transcriptionally-defined subpopulations remains to be assessed. A variation of Patch-Seq^55^, whereby RNA from imaged neurons is extracted and sequenced by single cell RNA-seq (scRNA-seq), could be a potential strategy to assess whether neurons exhibiting correlated responses belong to transcriptionally-defined subtypes. By generating *GAL4* drivers using the markers identified by scRNA-seq, we could determine the locations of these neurons in the brain and the types of circuits into which they integrate. Further, these subtype-specific markers would allow us track neurons that exhibit correlated changes in lactate flux, thereby allowing us to ask whether they exhibit correlated changes in other metabolic parameters such [glucose], [pyruvate], ATP/ADP ratio, and NAD^+^/NADH ratio using optical sensors^1^.

## ACKNOWLEDGEMENTS

We thank the Bloomington *Drosophila* Stock Center for fly stocks and Dr. Gregory Macleod for the *UAS-Laconic* fly line^31^. Live-cell fluorescence microscopy and image analysis was performed at the Center for Advanced Microscopy, a Nikon Center of Excellence, in the Department of Integrative Biology & Pharmacology at McGovern Medical School, UTHealth Houston (RRID: SCR_025962). We are grateful to Aditya Singh for his invaluable suggestions regarding the use of machine learning algorithms. This work was supported by National Institutes of Health (NIH) grants RF1AG069076 and RF1AG072176 to K.V. and K01HL143111 to T.I.M.

## AUTHOR CONTRIBUTIONS

M.S.P., E.R., T.I.M. and K.V. designed the research; M.S.P., E.R., R.G. and K. Vo performed the experiments; M.S.P. and K.V. analyzed data and wrote the manuscript.

## Methods

### Resource Availability

### Lead contact

Further information and requests for resources and reagents should be directed to and will be fulfilled by the lead contact, Dr. Kartik Venkatachalam.

### Materials availability

This study did not generate new unique reagents.

### Data and code availability

1. All data reported in this paper will be shared by the lead contact upon request.
2. All original analysis code has been deposited and made publicly available (https://github.com/kvenkatachalam-lab/Price-et-al-2025).
3. Any additional information required to reanalyze the data reported in this paper is available from the lead contact upon request.

## Method Details

### Drosophila husbandry

All flies were reared at room temperature on standard fly food (per 1L of food: 95 g agar, 275 g Brewer’s yeast, 520 g cornmeal, 110 g sugar, 45 g propionic acid, and 36 g Tegosept). We obtained the *d42-GAL4*, a driver fly line for transgene expression in motor neurons^32,33^, from Bloomington *Drosophila* Stock Center and the *UAS-Laconic*^31^ transgenic line from the laboratory of Dr. Gregory Macleod. *UAS-Perceval^HR^* was described previously^37^.

### Dissociation and culturing of *Drosophila* neurons

We dissociated neurons from *Drosophila* larval brains as described previously^7,37^. Briefly, we sterilized the cuticles of wandering 3^rd^ instar *d42>Laconic* larvae by immersion in ethanol, followed by a wash in sterile water. We dissected brains from these larvae in primary neuron culture media comprised of Schneider’s insect medium (S0146; Sigma-Aldrich) supplemented with 10% fetal bovine serum (FBS), antibiotic/antimycotic solution (A5955; Sigma-Aldrich), and 50 mg/ml of insulin (I6634; Sigma-Aldrich). After dissection, we washed the brains using fresh primary neuron culture media and transferred them to a dissociation solution comprised of filter-sterilized HL-3 (70 mM NaCl, 5 mM KCl, 1 mM CaCl_2_, 20 mM MgCl_2_, 10 mM NaHCO_3_, 115 mM sucrose, 5 mM trehalose, and 5 mM HEPES) supplemented with 0.4 mM L-cysteine (2430; Calbiochem), and 5 U/ml papain (P4762; Sigma-Aldrich). Tissues were then placed in an incubator at 25°C for 30 minutes to be enzymatically digested by papain. Following another wash in primary neuron culture media, we transferred papain-treated brains to a 1.5-ml tube containing 150 μl of primary neuron culture media for dissociation by trituration. We plated the dissociated neurons on 35-mm glass bottom dishes (D35-10-0-N; Cellvis) pretreated with concanavalin-A (C2010; Sigma-Aldrich). Cells were thereafter stored in a humidified container for four days in primary neuron culture media in a 25°C incubator. We washed the preparations with PBS once daily to eliminate any debris that may have remained from dissociation and to remove any yeast contamination.

### Confocal imaging of dissociated *Drosophila* neurons

Confocal imaging of dissociated neurons was performed as described^7,22,31,37^. Briefly, we used a Nikon A1R laser-scanning confocal microscope equipped with Nikon 40x/1.3 NA Plan-Fluor oil immersion objective and the “Perfect Focus System” for maintenance of focus over time. To assess lactate and glucose dynamics, we excited *Laconic*-or *FLII12Pglu-700μδ6*-expressing neurons with the 445 nm laser line and recorded emission signals at 488 nm and 535 nm. For ATP dynamics, we excited PercevalHR-expressing neurons with the 405 nm and 488 nm laser lines and collected at 535 nm (Venus). For pH correction, we excited the pHrodo Red AM ester with the 561 nm laser line and collected at 585 nm simultaneously multiplexed with either the ratiometric Laconic lactate or PercevalHR ATP biosensors. We imaged ∼20 cells per field and conducted at least 3 biological replicates for each treatment over the course of multiple days.

For live-imaging with agitation, plates of dissociated *d42>Laconic* neurons were filled with 3 mL of HL3 imaging media at room temperature prior to imaging. A 5 mm octahedral stir bar was placed near the rim of the plate. A magnetic mini-stirrer affixed upside-down to a ring stand was placed in close proximity above the stir bar. The lowest setting that induced rotation of the stir bar was used to agitate the mixture. The mini-stirrer was activated for 6 minutes prior to imaging and then left activated for the duration of the experiment.

For live-imaging without continuous agitation, we treated plates of dissociated neurons with the Invitrogen pHrodo Red AM pH indicator dye (P35372; ThermoFisher). Briefly, neurons were incubated for 30-minutes in 100 μl of a solution containing a 1x-dilution of pHrodo AM ester in primary neuron culture media. At the end of the incubation period, the pHrodo Red AM ester and culture media solution was replaced with primary neuron culture media. Primary neuron culture media was replaced with 100 μl of HL-3 at room temperature 10-minutes prior to imaging. Another ∼ 5-minutes taken to identify *Laconic*-expressing neurons ensured that cells had ∼15-minutes prior to the commencement of imaging.

We recorded baseline signals for 2 minutes prior to the application of 20 μl of HL-3 (mixing) or 20 μl of HL-3 containing metabolites or drugs at 6x final concentrations. Treatments were administered using an electronic pipette (Eppendorf Repeater E3) set at its 2^nd^-slowest dispense setting (#2) and a 100 μl Combitip. At the end of the 6-minute control or experimental conditions, we replaced the solutions with fresh HL-3 for 2 minutes. Subsequently, we applied 20 μl of a solution containing a 6x concentration of NH_4_Cl (10 mM final) in HL3 for 3-minutes to assess intracellular pH. This solution was then replaced with fresh HL-3 for an additional 3-minutes.

To examine neuronal viability, we treated plates of dissociated *d42>Laconic* neuron cultures with the Invitrogen LIVE/DEAD Fixable Near IR Dead Cell Stain (L34992; ThermoFisher). Prior to labelling, cells were washed twice with PBS and incubated for 30-minutes in 1x LIVE/DEAD Fixable Dead Cell Stain. As positive control, we treated cells with 0.1% (w/v) saponin (102855; MP Biomedicals) dissolved in primary neuron culture medium. After labeling, dissociated neuron cultures were fixed using 4% (w/v) paraformaldehyde (PFA) (15710; Electron Microscopy Sciences) in PBS solution for 15-minutes, followed by three 5-minute washes with 0.1% triton X-100 (w/v) in PBS. Neurons were mounted in DAPI Flouromount-G (0100-20; SouthernBiotech) and imaged on the Nikon A1R confocal microscope. We imaged multiple fields per dish using at least two dishes per saponin concentration.

To examine the distribution of glia, we fixed dissociated *d42>Laconic* neuron cultures with paraformaldehyde, followed overnight incubation at 4°C with mouse 8D12 anti-Repo antibodies (Developmental Studies Hybridoma Bank (DSHB)) along with 5% normal donkey serum blocking solution (D9663-10ML; SigmaAldrich). After washing, we incubated the neurons with AlexaFlour^TM^ 568 anti-mouse secondary antibodies (A11004; Invitrogen) for 1-hour at room temperature. Cells were mounted in DAPI Flouromount-G (0100-20; SouthernBiotech) prior to imaging on a Nikon A1R confocal microscope.

### Calculation of Laconic, PercevalHR, and FLII12Pglu-700**μδ**6 ratios and pH correction

For each field of view, we obtained fluorescence emissions from regions of interest (ROI) corresponding to individual neuronal cell bodies. To correct for background fluorescence, we subtracted the intensities of an ROI lacking cells from the intensities of the neuronal ROIs. Next, we corrected the emission intensities for fluorescence bleach, and then calculated the Laconic, PercevalHR, and FLII12Pglu-700μδ6 ratios with the bleach-corrected intensities (mTFP/Venus for Laconic, mCitrine/eCFP for FLII12Pglu-700μδ6, or 488 nm/405 nm for PercevalHR). For the long traces without buffer mixing (Figures 1B-1D and S1A), we normalized the Laconic and PercevalHR ratios to the starting values. For traces with buffer mixing at 2 minutes, we normalized the ratios to the means of their corresponding premixing values (i.e., baseline values).

We utilized an empirical pH correction protocol that first normalized the Laconic/PercevalHR ratios to the exponential fits of the premixing baselines in order to set the average premixing baselines to 1. NH_4_Cl-induced deviations of the pHrodo and Laconic/PercevalHR signals from baseline exhibited linear relationships, which were used to empirically correct for the changes in sensor ratios that stem solely from changes in cytosolic pH. We applied NH_4_Cl pulses at the end of each recording.

We determined the integrated changes in normalized, pH-corrected ratios by calculating the area under the curve (AUC) for each trace for 3 minutes from the time of mixing/treatment. We used the CausalImpact package^56^ in R to extrapolate the baseline, and thereby, estimated the ratios had we not applied the mixing/treatment. Difference between the AUC value of a trace and that of its extrapolated baseline, adjusted for the length of time after the treatment, represented the integrated change in ratios.

### Kinetic modeling of cellular metabolism

We developed simple mathematical models to simulate the dynamics of trehalose and lactate metabolism as dictated by unstirred layer effects, and rates of glycolysis and OXPHOS. The models were implemented in R using the deSolve package^57^ for solving the system of ordinary differential equations (ODEs). The system of ODEs was solved using the LSODA (Livermore Solver for Ordinary Differential Equations) algorithm implemented in the deSolve package. The metabolic system was described by coupled differential equations representing the temporal evolution of metabolites.

#### 1. Trehalose dynamics

d[trehalose]/dt = -v_trehalose_consumption

where *v_trehalose_consumption = k_trehalose * [trehalose]*

Here, *k_trehalose* is the rate constant for trehalose breakdown.

#### 2. Glucose dynamics

*d[glucose]/dt = -v_glucose_consumption + (2 * v_trehalose_consumption)*

where *v_glucose_consumption = k_glucose * [glucose] * (K_atp / (K_atp + [ATP])) * oxygen_factor_glucose*

Here, *k_glucose* is the rate constant for glucose consumption. We accounted for each trehalose molecule generating two glucose molecules. The velocity of glucose consumption was set to be contingent on demand for ATP using *(K_atp / (K_atp + [ATP]))* where *K_atp* is the half-maximal concentration of ATP. Since OXPHOS was implicit in the model, we also included a factor (*oxygen_factor_glucose*) to account for the complete oxidation of glucose. Oxygen-dependent regulation of glucose consumption was defined through a Hill equation: *oxygen_factor_glucose = [oxygen]^n_glucose / (K_oxygen_glucose^n_glucose + [oxygen]^n_glucose)*, where *K_oxygen_glucose* is the half-maximal concentration of oxygen needed for complete oxidation of glucose and (*n_glucose* = 2) was the Hill coefficient representing cooperative regulation of glucose consumption.

#### 3. Lactate dynamics

*d[lactate]/dt = -v_lactate_consumption + (lactate_coupling_factor * v_glucose_consumption)*

where *v_lactate_consumption = k_lactate * [lactate] * (K_atp / (K_atp + [ATP])) * (K_glucose / (K_glucose + [glucose])) * oxygen_factor_lactate*

Here, *k_lactate* is the rate constant for lactate consumption. We set the velocity of glucose consumption to be contingent on demand for ATP using *(K_atp / (K_atp + [ATP]))*. Further, *(K_glucose / (K_glucose + [glucose]))* where *K_glucose* is the half maximal concentration of glucose ensured that lactate consumption was contingent on glucose availability, i.e., lactate oxidation was driven by decreasing glucose levels.

Since OXPHOS was implicit in the model, we also included a factor (*oxygen_factor_lactate*) to account for the complete oxidation of lactate. Oxygen-dependent regulation of lactate consumption was defined through a Hill equation — *oxygen_factor_lactate = [oxygen]^n_lactate / (K_oxygen_lactate^n_lactate + [oxygen]^n_lactate)*, where *K_oxygen_lactate* is the half-maximal concentration of oxygen needed for complete oxidation of lactate and (*n_lactate* = 2) was the Hill coefficient representing cooperative regulation of lactate consumption.

#### 4. ATP homeostasis

d[ATP]/dt = v_atp_production -v_atp_consumption = 0

where *v_atp_production = v_atp_production_glucose + v_atp_production_lactate*; *v_atp_production_glucose = yield_atp_glucose * v_glucose_consumption*; and *v_atp_production_lactate = yield_atp_lactate * v_lactate_consumption*

Here, *yield_atp_glucose* and *yield_atp_lactate* refer to the ATP yield associated with complete oxidation of glucose and lactate, respectively. The rate of change in [ATP] was set to 0 to ensure that velocity of ATP production was equal to the velocity of ATP consumption.

#### 5. Oxygen utilization

d[oxygen]/dt = -v_oxygen_consumption

*where v_oxygen_consumption = k_oxygen_consumption * (v_glucose_consumption + v_lactate_consumption)*

Here, *k_oxygen_consumption* is the rate constant for oxygen consumption.

Parameters chosen for the models included — trehalose consumption (*k_trehalose* = 0.01); glucose consumption (*k_glucose* = 0.1); lactate consumption (*k_lactate* = 0.03); ATP from complete oxidation of glucose involving glycolysis, TCA cycle and ETC (*yield_atp_glucose* = 32); ATP from complete oxidation of lactate involving TCA cycle and ETC (*yield_atp_lactate* = 30); half-maximal concentrations (*K_atp* = 100, *K_glucose* = 10, *K_oxygen_lactate* = *K_oxygen_glucose* = 50); rate of oxygen consumption (*k_oxygen_consumption* = varied between 0.1 and 0.6 to ensure variable oxygen consumption rates); ATP demand (*[ATP]* = varied between 1 and 100 to ensure variable ATP demand).

The simulation was conducted in three sequential phases:

<u>1. Initial Phase (0–100-time units).</u> This phase represented the time equivalent to that prior to placing the dishes on the scope. Initial conditions were *[trehalose]* = 100, *[glucose]* = 10, *[lactate]* = 80, *[oxygen]* = 100.
<u>2. Intermediate Phase (100–200-time units).</u> This phase represented the time equivalent to the start of imaging to that equivalent to mechanical disruption of the unstirred layer. Initial conditions were end-state values from the first phase.
<u>3. Perturbation Phase (200–300-time units).</u> This phase represented the time equivalent to after mechanical disruption of the unstirred layer, i.e., when trehalose and oxygen concentrations were restored. Initial conditions were *[trehalose]* = 100, *[glucose]* = end-state values from intermediate phase, *[lactate]* = end-state values from intermediate phase, *[oxygen]* = 100.

### Jenks natural breaks classification

Using the classInt (https://cran.r-project.org/web/packages/classInt/index.html) R package, we implemented the Jenks Natural Breaks algorithm for stratification of the rate constants of lactate consumption. The classification process consisted of the following steps. First, we sorted the rate constants (k values) in descending order. Next, the dataset was partitioned into three distinct categories using the Jenks Natural Breaks optimization method, which minimizes within-class variance while maximizing between-class variance. This resulted in three categories. k values greater than or equal to the minimum threshold of the “High” category were assigned to cluster 1, those falling between the “Medium” category thresholds were assigned to cluster 2, and those below the minimum threshold of the “Medium” category were assigned to cluster 3.

### Neural network to predict patterns of changes in Laconic ratios

To predict the neurons’ response categories from baseline fluctuations in Laconic ratios prior to treatment, we trained a neural network on z-scaled Laconic time series data. The training model was comprised of a single hidden layer of 50 units with linear activation function. Units of the output layer also used the linear activation function. We implemented dropout regularization (rate=0.2) and L1/L2 regularization (rates=0.01) in the hidden layer to mitigate overfitting^44,45^. We used 5-fold cross-validation to predict post-mixing response. We determined prediction accuracies from 5-fold cross-validation using MSE as the metric of accuracy.

